# Evolutionary analyses of gene expression divergence in *Panicum hallii*: exploring constitutive and plastic responses using reciprocal transplants

**DOI:** 10.1101/2023.05.19.541545

**Authors:** Govinal Badiger Bhaskara, Taslima Haque, Jason E Bonnette, Joseph D Napier, Diane Bauer, Jeremy Schmutz, Thomas E Juenger

## Abstract

The evolution of gene expression is thought to be an important mechanism of local adaptation and ecological speciation. Gene expression divergence occurs through the evolution of cis-polymorphisms and through more widespread effects driven by trans-regulatory factors. Lovell et al. (2018) studied expression divergence between two ecotypes of *Panicum hallii* using expression quantitative trait loci (eQTL) analyses and discovered a pre-dominance of cis and several trans-regulatory divergences. Here, we explore expression and sequence divergence in a large sample of *P. hallii* accessions encompassing the species range using a reciprocal transplantation experiment. We observed widespread genotype and transplant site drivers of expression divergence, with a limited number of genes exhibited genotype-by-site interactions. We used a modified F_st_-Q_st_ outlier approach (*Q_PC_* analysis) to detect local adaptation. We identified 514 genes with constitutive expression divergence above and beyond the levels expected under neutral processes. However, no plastic expression responses met our multiple testing correction as *Q_PC_* outliers. Constitutive *Q_PC_* outlier genes were involved in a number of developmental processes and responses to abiotic environments. Leveraging the earlier eQTL results, we found a strong enrichment of expression divergence, including for *Q_PC_* outliers, in genes previously identified with cis and cis-drought interactions but found no patterns related to trans-factors. Population genetic analyses detected elevated sequence divergence (F_ST_, D_XY_) of promoters and coding sequence of constitutive expression outliers, but little evidence for positive selection on these proteins. Our results are consistent with a hypothesis of cis-regulatory divergence as a primary driver of expression divergence in *P. hallii*.

## Introduction

Gene expression variation plays a major role in phenotypic evolution, both within and between species, facilitating adaptation to their native habitats (Fay and Wittkopp, 2008; Gilad et al., 2006; Necsulea and Kaessmann, 2014; Nourmohammad et al., 2017; Price et al., 2022; Romero et al., 2012; Whitehead and Crawford, 2006). Gene expression is a dynamic trait that responds to environmental changes, leading to plasticity in morphological, physiological, and fitness traits. As populations adapt to changing environments, they can exhibit constitutive or plastic expression divergence. Constitutive expression divergence may occur when plasticity is costly or maladaptive, whereas plasticity can be favored when the environment is heterogeneous and requires multiple optimal responses throughout an organism’s life cycle or when the environmental cue is predictable (Hendry, 2015; Murren et al., 2015; Mäkinen et al., 2017; Li et al., 2019). When the environmental response is heritable and plasticity is beneficial, natural selection can reinforce it leading to evolution of plastic expression (Campbell-Staton et al., 2021; Crispo 2007). This adaptive expression plasticity can play a major role in enhancing resilience to climate change (Nicotra et al., 2010; Brooker et al., 2022). However, there is currently a lack of empirical research that investigates adaptive constitutive and plastic expression divergence in natural populations and their underlying regulatory architectures.

Both *cis*- and *trans*-acting regulatory architectures contribute significantly to expression variation (Fay and Wittkopp, 2008). The relative contributions of cis- and trans-variants differs depending on the evolutionary timeline, with inter-species variation occurring over longer periods and intra-species variation occurring over shorter periods (Stern and Orgogozo, 2008). Cis-regulatory elements (CREs) are modular and allow expression changes specific to tissues, life stages or environmental conditions therefore are more likely to have additive effect for fitness than trans-acting regulatory mutations which influence gene regulatory networks with wider connectivity, and tend to have larger pleiotropic effects (Prud’homme et al., 2007; Metzger et al., 2017; Vande Zande et al., 2022). This has led to the hypothesis that cis-variants accumulate over time since they possess fewer selective constraints due to less negative pleiotropy as compared to trans-variants (Prud’homme et al., 2007). In yeast, flies and Arabidopsis cis-variants contribute more to inter-species differences in gene expression than trans-variants (Ronald et al., 2005; Wittkopp et al., 2008; Tirosh et al., 2009; Shi et al., 2012), while in maize and *Panicum hallii* (a perennial grass), cis-variants account for a greater proportion of intra-species gene expression variation (Schadt et al., 2003; Guo et al., 2004; Springer and Stupar, 2007; Lovell et al., 2016; Lovell et al., 2018). Overall, the relative contributions of cis- and trans-variants within and between species vary considerably across different model systems. This variability may be attributed to factors such as genetic diversity, selective pressure, divergence time, environmental heterogeneity and even the methods used to estimate expression variation (Li and Burmeister, 2005; Meiklejohn et al., 2014). Nevertheless, given the pleotropic constraints, it is suggested that purifying selection may strongly shape trans-variants, while positive selection contributes significantly to cis-expression variation leading to adaptive divergence (Schaefke et al., 2013).

Divergence in gene expression has been suggested to correlate with the evolutionary rate of proteins (Ingvarsson, 2007; Slotte et al., 2011). However, the relationship has been the subject of much debate due to contrasting results between studies. For example, Renaut et al (2012) and Moyers and Rieseberg (2013) have found no significant relationship between these two factors. Additionally, the types or direction of selection driving this association also varies between studies (Hodgins et al., 2016). In Drosophila, positive selection has been proposed as a factor driving the association between expression divergence and sequence evolution, while in mammals and pines, the weak association between expression divergence and sequence evolution has been attributed to relaxed purifying selection and genetic drift (Nuzhdin et al., 2004; Liao and Zhang, 2006; Hodgins et al., 2016).The reasons for the variability remain unclear, but differences in sequence evolution, the efficacy of selection, and the timescale of evolution are likely to contribute to the observed patterns.

Perennial grasses provide a unique opportunity to study local adaptation. We developed a diploid C4 perennial bunchgrass*, Panicum hallii,* as a model system to study ecology, genomics and molecular evolution (Lowry et al., 2013; Lovell et al., 2018; Palacio-Mejía et al., 2021). Gene expression changes have been reported in its adaptation to abiotic stresses and local environments (Lovell et al., 2016; Lovell et al., 2018; Haque et al., 2022). Notably, the expression of drought responsive genes between inland and coastal ecotypes of *P. hallii* was influenced by CREs (Lovell et al., 2016) and over 50% of transcripts in *P. hallii* exhibited cis expression quantitative loci (eQTL) underlying expression divergence between ecotypes, indicating the prevalence of cis-expression divergence (Lovell et al., 2018).

In this study, we conducted a field reciprocal transplant experiment using natural genotypes of *P. hallii*. Our objectives were to: i) detect gene expression variation resulting from genetic differences or plastic responses to the environment, along with their interactions, ii) dissect the heritable constitutive and plastic components of gene expression variation using an outlier-based testing framework (Josephs et al., 2019) designed to detect divergent selection on gene expression, and iii) evaluate the relationship between constitutive gene expression divergence and the rate of sequence and protein evolution. We identified hundreds of genes with constitutive expression divergence beyond neutral expectation, but we detected a very weak signal for adaptive plastic divergence. Moreover, our results suggest that the genes exhibiting constitutive expression divergence in our study were not under positive selection but evolve faster compared to other background genes due to relaxed purifying selection.

## Results

### Population genetic analysis of *Panicum hallii*

Natural populations of *Panicum hallii* demonstrate substantial divergence in their morphology and physiology (Lowry et al, 2013) which could be due to historical demographic process and/or adaptation to ecologically different native habitats. To study the role of gene expression divergence in adaptive differentiation we performed a field reciprocal transplant experiment on 86 genotypes sampled from the natural range of coastal and inland habitats (Supplementary Table S1). Discriminant analysis of principal components (DAPC) on a random set of 50,000 putatively neutral SNPs from the sampled population inferred two major genetic clusters that were consistent with two previously described ecotypes (*var. hallii* and *var. filipes*) (Figure 1A, Lovell et al., 2018). *var. hallii* is distributed over a much wider inland geographic area from Texas to Arizona (hereafter referred to as inland cluster, Figure 1B, dark blue and red points), while *var. filipes* is restricted to the southeastern portion of Texas (hereafter referred to as the coastal cluster, Figure 1B, light blue points). Given the small sample size of coastal cluster (n=11) from a relatively limited geographic area, we chose to focus subsequent hierarchical cluster analysis on the inland population only. Bayesian clustering analysis implemented in STRUCTURE supports two sub-clusters of the inland collections (K=2) according to the Δk method (Supplementary Figure S1, Supplementary Table S1). We designated these two clusters as the ‘Northern’ (n=53) and ‘Central’ (n=22) clusters. The Northern cluster was found from Central Texas to Arizona spanning a steep precipitation gradient (Figure 1A, dark blue points), while the Central cluster is primarily located in Central Texas (Figure 1A, dark red points). These three genetic clusters were congruent with the major clusters found in earlier studies of *P. hallii* natural accessions (Lovell et al, 2018, Palacio-Mejía, 2021). However, Lovell et al (2018) and Palacio-Mejía et al (2021) found finer groups within each cluster likely as a result of their denser sampling. Lovell et al (2018) estimated the divergence time of the inland and coastal populations of *P. hallii* as over a million year. In order to polarize the ancestry between inland and coastal genetic clusters we constructed a phylogenetic tree from the concatenated alignment of the one-to-one orthologs among one inland and one coastal *P. hallii* genome reference along with *P. virgatum* and *Setaria viridis.* The ancestral state was inferred using *P. virgatum* and *Setaria* as outgroups. We reasoned that the more ancestral ecotype would contain a greater proportion of ancestral allele states, and use this framework to test for the order of divergence. However, we found equivocal results as the proportion of ancestral alleles to derived alleles was roughly equal for both inland and coastal populations (95% bootstrapped confidence interval for the probability that coastal genetic cluster is ancestral= 0.47-0.49).

**Fig. 1.**
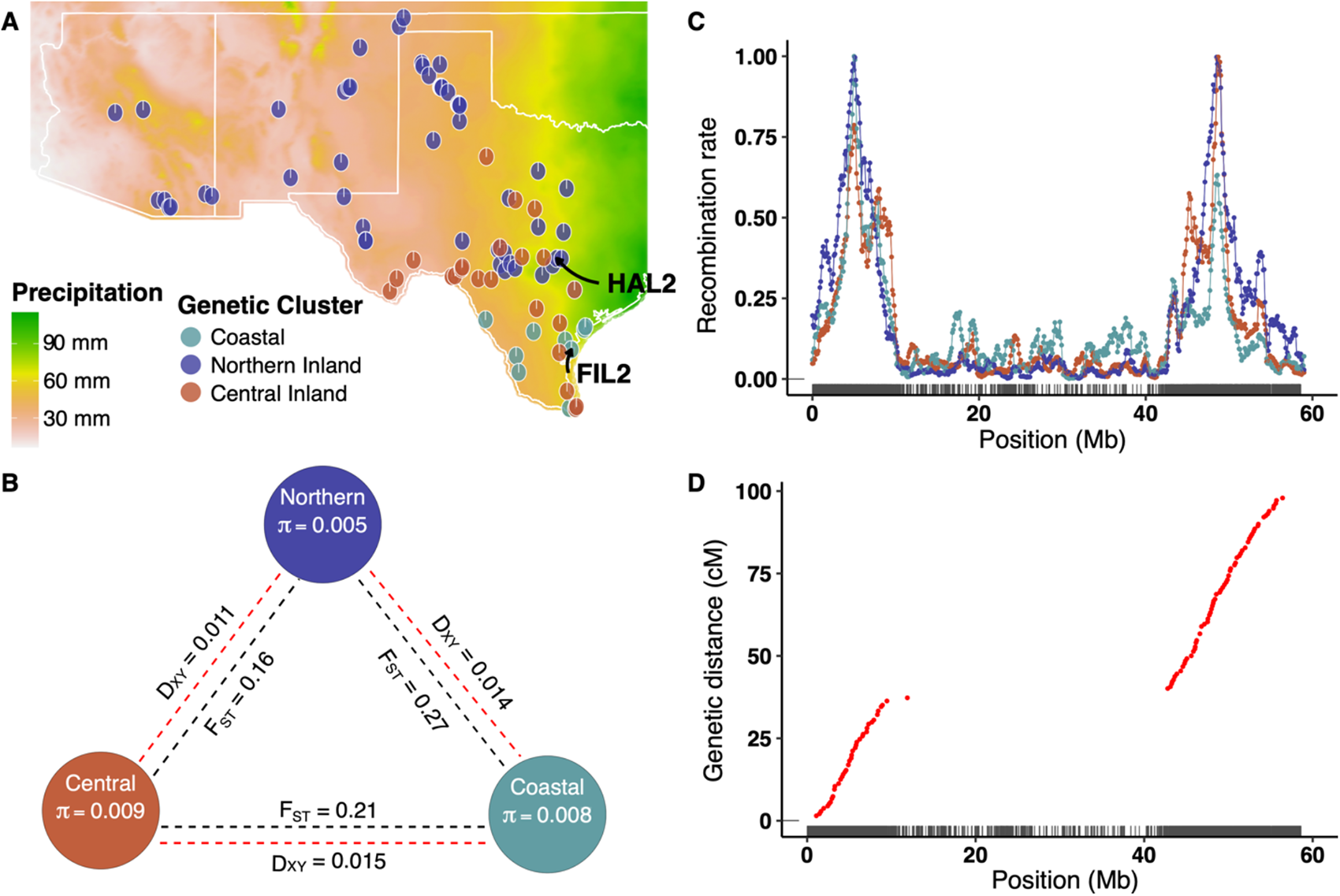
Genetic analysis identified three major genetic clusters of *P. hallii*. A. Genetic structure analysis and geographical distribution of *P. hallii* populations. The background Raster plot was constructed from the average annual precipitation (BIO12) of 2.5-minute spatial resolution (http://worldclim.org/version2). Points represents the origin of collections for each accession. Point colors represent genetic subpopulations. Two major genetic cluster (Coastal and Inland) were detected by Discriminant Analysis of Principal Component (DAPC) of 86 accessions of *P. hallii*. Subsequent Bayesian hierarchical cluster analysis (STRUCTURE) inferred two genetic clusters within the inland population (Northern Inland: dark blue, Central Inland: brick red). A pie in the plot represents the proportion of posterior probability for each group by STRUCTURE (K=2). Coastal populations (light blue) were plotted based on the origin of collection site without ancestral population inference. B. Genome-wide Nucleotide diversity within genetic cluster and (π) and Divergence statistics (FST, D_XY_) between genetic clusters. C. Recombination landscape of individual genetic cluster across chromosome (Chromosome 1 as an example). Population-scaled recombination rate was plotted on y-axis and x-axis represent the physical position of the chromosome. The rug at the bottom of the figure represents gene density. D. Genetic map of *P. hallii* from a Recombinant Inbred Line population (chromosome 1 as an example). The x-axis represents the physical distance and y-axis represents genetic distance. The gap between the arms of chromosome 1 depicts the heterochromatin region which demonstrates low recombination rate (panel C).

To investigate genetic differentiation within *P. hallii*, we estimated relative (F_ST_) and an absolute measure of divergence (D_XY_) between genetic clusters and nucleotide diversity (π) within each cluster at 20-Kb non-overlapping windows genome-wide. Relative diversity varied significantly (*P* < 0.05) among all three possible pairwise contrasts of genetic clusters (Fig. 1B). The highest F_ST_ was observed in Coastal-Northern contrast (mean F_ST_=0.27), followed by the Coastal-Central contrast (F_ST_=0.21) with the least relative divergence observed in the Central-Northern comparisons (F_ST_=0.16). Similar to relative diversity, the absolute measure of divergence among genetic clusters was significantly different for all three pairwise comparisons (*P* < 0.05). However, here the Coastal-Central contrast revealed the greatest divergence (mean D_XY_=0.015) followed by the Coastal-Northern contrast (mean D_XY_=0.014), and finally the Central-Northern contrast (mean D_XY_=0.011). The highest nucleotide diversity was observed in the Central cluster (mean π=0.009) followed by Coastal (mean π=0.008) and Northern clusters (mean π=0.005). Overall, we observed Coastal vs. Inland pairs were more diverged compared to inland clusters and the Northern cluster had relatively low nucleotide diversity than the other genetic clusters.

Recombination rate has been reported to vary between populations and along chromosomes in many plant species (Dreissig et al, 2019, Schreiber et al, 2022) and can be a major driver of molecular evolution (Dapper and Payseur, 2017). In particular, regions of low recombination are often associated with reduced genetic diversity and reduced effectiveness of natural selection. We estimated the mean scaled recombination rate (ρ) for each genetic cluster based on population genetic analyses of whole genome resequencing data. We found significant difference in ρ between genetic clusters of *P. hallii* (post hoc adjusted *P* < 0.05): the highest ρ was estimated for Northern genetic cluster (ρ=0.33/kb) followed by Central (ρ=0.24/kb) and Coastal genetic cluster (ρ=0.2/kb). Noticeably, we observed an elevated recombination rate around euchromatin regions (near the distal end of the chromosome) compared to heterochromatin regions near the pericentromeric regions (Pearson’s correlation coefficient of population-scaled recombination rate and gene density for Coastal, Central and Northern genetic cluster were 0.4, 0.5 and 0.7 respectively, *P* < 2.2e^-16^) (Figure 1C; Supplementary Figure S2). This landscape of recombination was congruent with the number of crossover events estimated from a Recombinant Inbred Line (RIL) population (HAL2 x FIL2; Khasanova et al. 2019) generated from Northern and Coastal reference parents (Spearman’s Rank correlation coefficient = 0.5; *P* < 2.2e^-16)^ (Figure 1D). The highly recombinant region encompassed almost half (46%) of the genome of *P. hallii* (Supplementary Table S2) and contains that vast majority of coding genes (82%; 27,271). Nevertheless, 5,992 genes were found in the largely pericentromeric regions.

### Plant water status and transcriptome changes in response to genetic variation and transplant sites

We used a field reciprocal transplant experiment to study the physiology and gene expression variability among different genetic clusters grown in native and foreign habitats of origin (Figure 2A). We assessed plant water status by measuring leaf relative water content (RWC) at the time of tissue harvest for transcriptome sampling. RWC indicates plant water status in terms of the hydration state of plant and it can be a strong driver of plant performances and gene expression changes (Jones et al., 2006, Des Marais 2013). We analyzed RWC for the effect of Genetic clusters (G), field environment (E), and their interaction (GxE). In this framework, we interpret a significant main effects of G as constitutive divergence in RWC, a significant effect of E as plastic responses of RWC to planting location, and a significant interaction as genetic variation in the plastic response of genotypes to planting sites in terms of water status. We observed a significant G (F_(df=2)_ = 4.032, *P <* 0.05), E (F_(df=1)_ = 147.3, *P* < 0.001), and GxE (F_(df=2)_ = 6.9, *P <* 0.01) for RWC (Figure 2B) revealing genetic variation in whole plant physiology in response to our transplant locations. Overall, the three genetic clusters diverge in RWC at the coastal site (F_(df=2)_ = 8.184, *P <* 0.001), and differ little at the inland site (F_(df=2)_ = 0.7, *P >* 0.1). At the coastal site, the native coastal cluster maintained the highest RWC, followed by Northern and Central clusters.

**Fig. 2.**
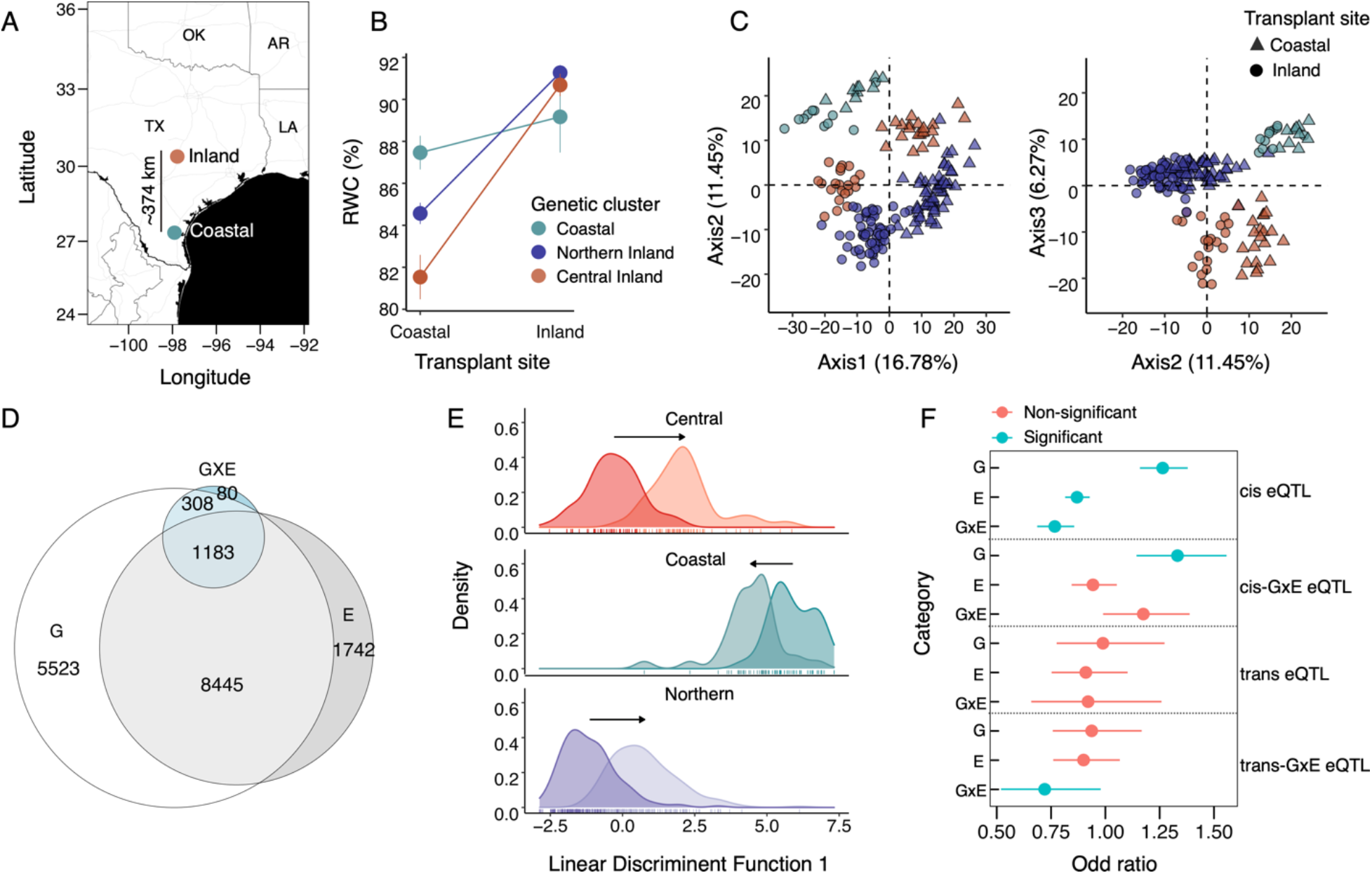
Effects of population and transplant site on gene expression. A. Map of study sites at the Central (Inland) and South Texas (Coastal). B. Reactions norms of Relative Water Content (RWC) among population in response to transplant sites. Colored symbols represent the genetic clusters identified from SNPs. C. Principal Component Analysis (PCA) of gene expression variation for the top 1000 most variable genes. Colored symbols represent the genetic cluster as shown in B. D. Venn diagrams showing gene expression distribution for genotype (G), field environment (E), and genotype-by-environment interactions (G×E) for given population contrasts. E. Discriminant analysis of principle components (DAPC) on total transcriptomic space representing the effect of origin/genetic cluster on expression plasticity. The x-axis represents the linear discriminant function in multivariate space along which the difference between native habitat to transplant site is maximized. Each panel depict the distributions of DAPC score of focal genetic cluster at native site (darker shade) and at transplant site (lighter shade). The distance between the mean of a focal genetic cluster at home to the mean at transplant site was marked by arrow which represents transcriptomic plasticity. Plasticity was significantly different for all three pairwise comparisons (adjusted p-value < 0.05).

Similar to plant water status, transcriptomic changes can be responsive to environmental shifts due to transplantation to foreign habitat. The global patterns of expression variation based on a principal component analysis (PCA) detected clear expression differentiation among field environments and *P. hallii* populations (Figure 2C). PCA axis 1 (explaining 16.78% variance) was associated with a strong environment effect, while axis 2 (11.45% variance) and 3 (6.27% variance) were related to population divergence in gene expression. Overall, variation in gene expression was in concordance with the three distinct genetic clusters of *P. hallii* detected from our population genetic analysis. As such, we first tested for the global effect of G, E and GxE on each expressed gene by factorial model which includes all three genetic clusters across two field environments. We found a strong transcriptomic response of genetic cluster (15,656 genes with significant G effect) and plasticity (12,387 genes with significant E effect), with many fewer genes exhibited genetic cluster dependent plastic response across transplant sites (1,877 genes with significant GxE effect) (Figure 2D, Supplemental Table S3). Similarly, GxE explained a much lower proportion of variance in gene expression than G or E (Figure 2D, Supplementary Figure S3). Noticeably, we observed enrichment of G category genes and depletion of GxE category genes at low recombining regions of the genome compared to the highly recombining regions (P-value < <0.05). However, no significant enrichment was found for plastic genes (E).

To further understand how the global patterns of gene expression plasticity varies among different genetic clusters, we analyzed genome wide expression profile in multivariate space by implementing Discriminant Analysis of Principle Components (DAPC). The linear discriminate function was derived to maximize the variation between genetic clusters with respect to their native habitats (inland genotypes as inland site vs. coastal genotypes at coastal site) and transplants samples were projected on that discriminant axis to estimate expression plasticity of a focal genetic cluster (Figure 2E: DAPC plot). We observed that the magnitude of expression plasticity varied between different genetic clusters and each pairwise comparisons (significant at each pairwise comparison level, adjusted *P* < 0.05). The Coastal genetic cluster demonstrated the least plastic transcriptome response compared to any of the inland clusters. Subsequently, we conducted post hoc pairwise comparisons between genetic clusters and transplant sites on each expressed gene. Consistent with the global analysis, pairwise analysis that includes two genetic clusters at a time also showed strong G and E transcriptomic responses, and only a few hundred genes exhibited significant GxE effect (Supplemental Table S4, S5, S6). Similarly, G and E explained a relatively higher proportion of variance in gene expression than GxE for all three contrasts (Supplementary Figure S3).

### Genetic architecture of expression divergence

Previously, we studied expression divergence between two inland and coastal ecotypes of *P. hallii* using eQTL analyses in a field drought experiment (Lovell et al., 2018). In that study we discovered a pre-dominance of *cis*-regulatory divergence, along with several *trans*-QTL that were enriched for drought responsiveness. Here, we leveraged our more expansive study of inland and coastal expression diversity to test the generality of these results. In particular, we ask whether differentially expressed genes detected in the current study (G, E, or GxE) were enriched for the occurrence of inland/coastal eQTL (*cis*, *trans*, *cis*-eQTL x drought*, trans*-QTL x drought) (Figure 2F) from our earlier study. We observed an over-representation of *cis*-eQTL and *cis*-eQTL x drought in G genes, along with an over-representation of *cis*-eQTL x drought genes in our GxE lists. Interestingly, we observed a significant under-representation of *cis*-eQTL genes in our E and GxE categories and a significant under-representation of *trans*-QTL x drought genes in our GxE genes. Overall, we found more overlap among studies in terms of *cis*-and *cis*-eQTL x drought than for *trans* factors.

### Detecting selection on gene expression divergence from reciprocal transplantation

Given the considerable gene expression divergence present among *P. hallii* populations, we subsequently asked to what degree the observed divergence in gene expression and its plastic response to the environment is the result of neutral or adaptive evolutionary processes. To infer the past action of natural selection we implemented the *Q*_PC_ method. This method uses PCs of relatedness matrix to estimate additive genetic variance and test for departure from neutral expectations in trait divergence. Figure 3A demonstrates how the lower order PCs separate the major genetic clusters of *P. hallii* population across the axes on which the expression divergence was tested. The neutral expectation was derived from estimated additive genetic variance from higher order PCs. Figure 3B and 3C delineated examples of trait evolve neutrally and under selection (excessive divergence) respectively on the first PC which separated coastal genetic cluster from inland clusters. As mentioned above we noticed remarkably variability in the pattern of historical recombination across the *P. hallii* genome, with likely important consequence for evolutionary dynamics in the recombing and non-recombining portions. As such, we employed the *Q*_PC_ method in two different modes to obtain constitutive component of expression divergence: i) Relaxed mode where we applied *Q*_PC_ method on the mean expression of all expressed genes (18,773) across sites and ii) Constrained mode on the mean expression of genes which resided in the highly recombined regions (15,735) across sites. Similarly, we explored the plastic component of expression divergence by studying the difference in expression between field sites for each accession. We designate these patterns as constitutive divergent expression (CDE) and adaptive plastic divergent expression (PDE).

**Fig. 3.**
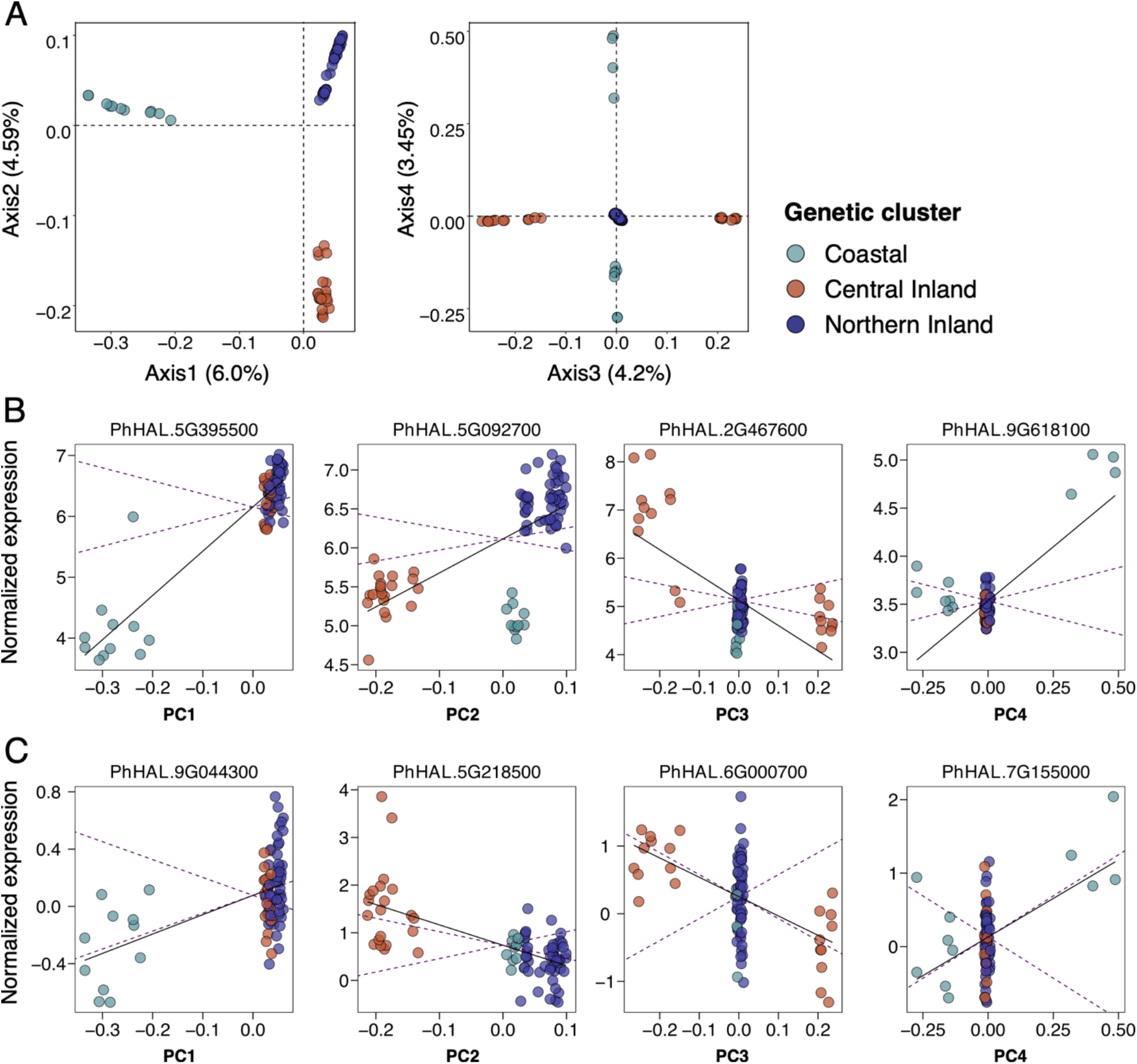
Outlier gene expression for reciprocal transplantation based on Q_PC_ analysis. A. The first four genetic PCs of the kinship matrix. Each circle represents different accession which are colored by three different genetic clusters. Each axis is labelled by the percentage of variation explained by that PC compared with the total variation. B. Genes that showed evidence of selection on their constitutive divergent expression (CDE). The mean-centered expression of the genes was plotted against the eigen value of corresponding PCs for selected genes that were identified as outliers from neutral expectation. Each point represents a single *P. hallii* natural accession and is colored by genetic clusters. The linear regression is plotted as a solid line and the 95% confidence intervals (CI) of the neutral expectation are shown as dotted lines in each plot – expression values outside of the CI envelope are taken as putative adaptive divergent expression. C. Genes that demonstrated plastic divergent expression (PDE) between inland and coastal sites. The data format is the same as described for B.

For CDE we found 1006 and 514 unique genes with excessive divergence beyond neutral expectation for the first five PCs by relaxed mode and constrained mode respectively (FDR threshold < 0.1). While implementing relaxed mode analysis, we observed an excess of genes with constitutively elevated expression divergence from the low recombined regions compared to the expressed background genes (Fisher’s exact test odd ratio 5.4; *P* < 2.2e^-16^). As such, from here onward we only considered constrained mode for both CDE and PDE. In the CDE category, PC1 accounted for almost half of these genes (224) (Supplemental Table S7). GO enrichment analysis of these genes identified ten GO terms including terms related to response to stimulus and ion binding (Supplemental Table S8; nominal P < 0.05). PC2 accounted for 161 genes with an enrichment of 19 GO terms such as pollen-pistil interaction and response to stress. PC3, PC4 and PC5 accounted respectively for 96, 50 and 65 genes which exhibited significant divergent expression. CDE genes on PC3, PC4 and PC5 were enriched with GO terms including terms related to anion binding, and endopeptidase activity.; cellular protein metabolic process, ATPase coupled transmembrane transporter; and RNA processing and chromosome segregation respectively. Figure 3B depicts the expression profile of a few example genes along the corresponding PCs for which non-neutral expression divergence has been inferred. For instance, PhHAL5G395500 (high affinity potassium transporter 1 demonstrated expression divergence along PC1 between coastal and inland populations (Supplementary Table S7). We have detected 82 genes that show evidence of selection on multiple PC indicating that adaptive expression divergence could be a function of divergence across more than two genetic clusters. For example, PhHAL.5G092700 (oxidoreductase family protein) demonstrated excessive expression divergence for both PC1 and PC2. PhHAL.2G467600 (AP2 domain containing protein) and PhHAL.9G618190 (heat shock protein 60) were plotted as examples of divergently expressed on PC3 and PC4 respectively (Figure 3B).

For PDE category, we failed to detect any statistically significant signal of non-neutral divergence that passing multiple test correction. However, we detected 29 genes that passed the nominal *p-value* threshold (< 0.05) (Supplementary Table S9). The majority of these were detected along PC2 (11 genes) and were related to plastic expression divergence between the Central to other two genetic clusters (Figure 3C). Only three genes were detected with significant divergence between the inland and coastal genetic clusters along PC1. Nine and four genes were detected along PC3 and PC4 between the central and northwestern inland genetic clusters. We plotted PhHAL.9G044300 (Galastosyl transferase family protein), PhHAL.5G218500 (chitinase A), PhHAL.6G000700 (cysteine proteinase superfamily protein) and PhHAL.7G155000 (ferric-chelate reductase) as examples of plastic expression divergence between populations along PC1, PC2, PC3, and PC4 respectively (Figure 3C).

Overall, we detected strong evidence for adaptive constitutive gene expression divergence (514 CDE genes) and weaker signal for adaptive plastic gene expression (PDE) divergence in our study. Moreover, we observed enrichment of both *cis* (odd ratio 1.7, Fisher’s Exact Test *P* =9.7e^-8^) and cis-GXE eQTL genes (odd ratio 1.7, Fisher’s Exact Test *P* =3.2e^-4^) but no enrichment of trans and trans GxE eQTL in our CDE category.

### Relationship between expression and sequence divergence

In order to understand the relationship between expression divergence and sequence evolution, we compared patterns of polymorphism and divergence between genes with putatively adaptive CDE and a set of background genes (15,237 expressed genes with non-significant divergence (NDE)). Given the relatedly weak signal for PDE genes we chose to exclude these from our analyses. Here, we looked at within and between genetic cluster diversity at the different annotated gene features including coding sequence (CDS), intron, 3′ and 5′-untranslated regions (UTR). To explore patterns of diversity and divergence in putative regulatory sequence, we also studied proximal promoter (2 Kb upstream from transcription start site) of target gene models. We observed significantly higher relative divergence (F_ST_) at CDE genes compare to NDE genes for all three possible pairwise contrasts at each gene feature separately (Figure 4A, Supplementary Table S10).

**Fig. 4.**
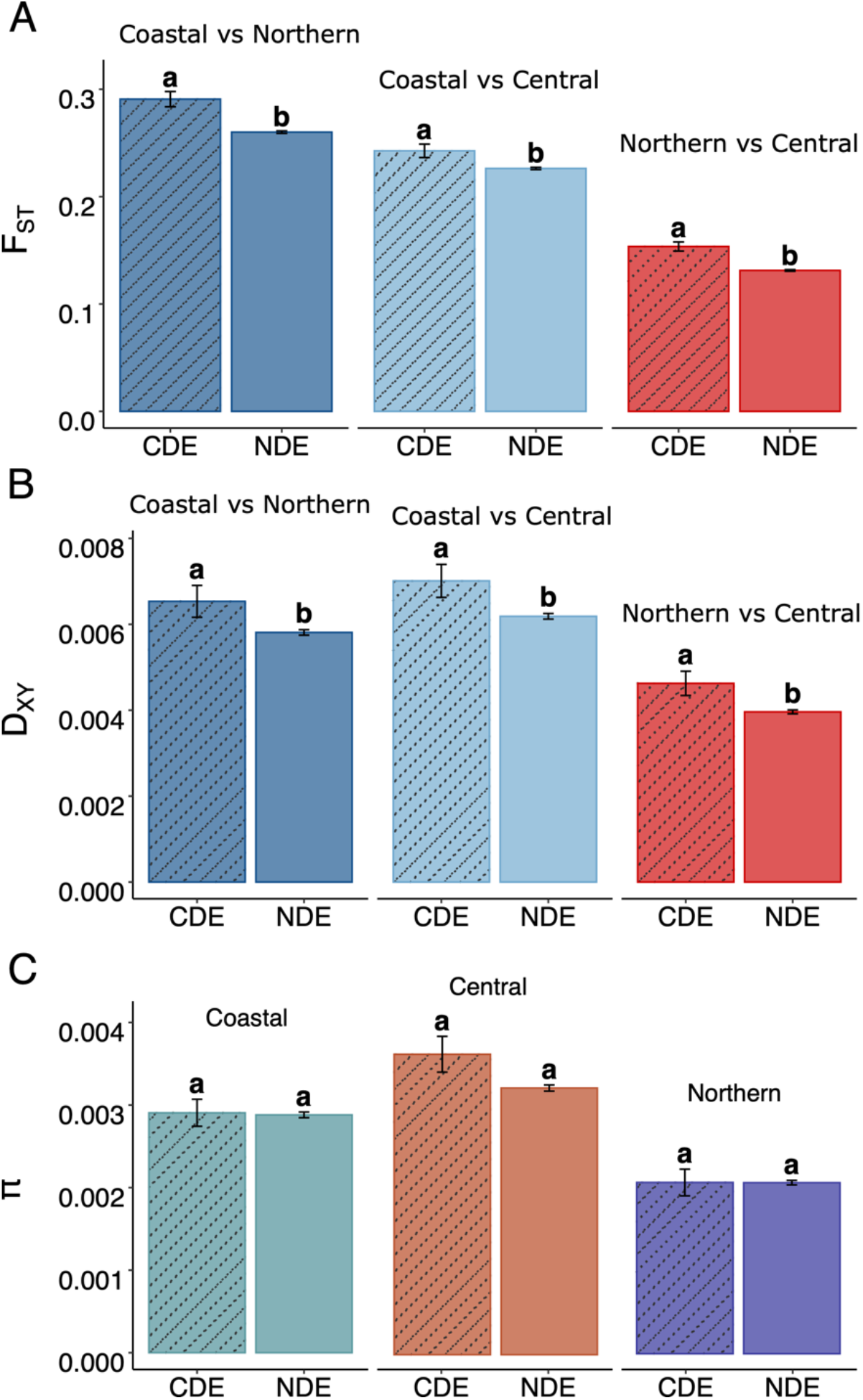
Divergence statistics at Coding Sequences (CDS) of constitutive divergent expression (CDE) and non-significant divergent expression (NSE) genes A) relative fixation index (F_ST_), B) absolute divergence (D_XY_) and C) nucleotide diversity (pi)

To address whether genetic variation between or within populations drive this pattern of differentiation, we estimated absolute divergence (D_XY_) and nucleotide diversity (π) statistics at each gene features. We observed significantly higher absolute divergence (D_XY_) for all three contrasts for CDE genes compared to NDE at CDS, 5′-UTR and promoter regions (except Northern vs. Central contrast at 5′-UTR and promoter) (Figure 4B, Supplemental Table S11). However, there was no difference in D_XY_ at intron and 3′-UTR between CDE and NDE category genes in any pairwise contrast. No significant difference in nucleotide diversity (π) within each genetic cluster was observed except for Northern and Central inland cluster at promoter regions (Figure 4C). Overall, we observed excessive relative and absolute divergence at CDS and promoter regions for CDE genes compare to NDE genes.

### Tests for positive selection for proteins with divergent expression

Since we observed higher relative and absolute sequence divergence among populations for the protein coding sequences of CDE genes, we hypothesized that the proteins of these genes might exhibit different rates of evolution among *P. hallii* lineages. Therefore, we quantified the ratio of substitution rates for non-synonymous and synonymous sites (dN/dS) of all one-to-one orthologous gene pairs of coastal and northern inland lineage based on existing annotated reference genomes. In total, we evaluated 184 CDE genes. We observed significantly higher evolutionary rates for CDE genes compare to NDE genes (Mann-Whitney U test *P* < 0.01). Down-sampling the NSE category to the number of CDE category also demonstrated higher evolutionary rate in CDE category (Mann-Whitney U test *P* < 0.01. We also detected a negative correlation between both dN/dS and dN and average expression level across site (p-value of Spearman’s Rank correlation test < 0.05). Therefore, we further tested for partial correlation of constitutive divergence expression with dN/dS and dN while controlling for mean expression and detected a positive correlation for both the factors (CDE genes had higher dN/dS and dN; p-value of Spearman’s Rank correlation test < 0.05). Similarly, mean expression level of CDE genes was lower than NDE genes (Mann-Whitney U test *P* < 0.01). However, we did not find any difference for the frequency of genes with dN/dS >1 in CDE group compare to NDE group (Chi-square test *P* = 0.66) which could be likely candidates for positive selection. Specifically, we detected 4 and 129 genes in CDE and NSE categories respectively with estimated dN/dS > 1.

Finally, to identify genes experiencing positive selection in the *Panicum* lineage we compared the likelihood of PAML’s (version 4.9) sites-specific selection model (M2a) with that of neutral model (M1a) across the *Panicum* phylogeny (Yang et al,2007). Here, we included the two sub-genomes (N and K genomes) of *Panicum virgatum* to infer ancestral states and restricted our analysis to 7,419 CDE genes with one-to-one ortholog pairs for all six pairwise comparisons. After correcting for multiple tests, we detected 163 genes which exhibited evidence of positive selection (Supplemental Table S12). However, we did not observe enrichment of CDE genes in this list of positive selection candidates (Chi-square test *P* = 0.66).

## Discussion

### Patterns of gene expression divergence in *P. halli* population

In this study, we decomposed the heritable component of constitutive and plastic transcriptome-wide gene expression divergence in *P. hallii* using a reciprocal transplant experiment at two native sites. In congruence to earlier studies (Whitehead and Crawford, 2006b; Staubach et al., 2010; Groen et al., 2020) we detected most of expressed genes were evolving (nearly) neutrally. Noticeably, we detected relatively strong signals of constitutive expression divergence beyond the expectation of neutral evolution for several hundred genes (514 genes) and only a weak signal of putatively adaptive plastic expression divergence. Constitutive expression divergence (Hodgins et al., 2016; Fuess et al., 2021) and population dependent plastic responses of transcript to changing environment or treatment (Kenkel and Matz, 2016; Mäkinen et al., 2017) have been reported in many study systems. However, heritable expression divergence can also result from random genetic drift, but only a handful of studies have implemented an analysis framework to incorporate a neutral model to detect the effect of natural selection. One example is a study by Whitehead and Crawford (2006a) on teleost fish populations that inferred that expression divergence of several genes in a common garden experiment was due to natural selection. Groen et al. (2020) estimated stronger selection on gene expression in drought conditions compared to wet conditions and reported that rice accessions which were more plastic experienced fitness benefits. Adaptive plastic expression response to changing environments can provide resilience to a population and therefore is of central interest, especially in the context of recent climate change. In this study, we observed a weak signal of such responses in *P. hallii*. Several factors such as high cost of plasticity, lack of genetic variation towards plastic response, low gene flow, relaxed selection due to environmental variability, or maladaptive plasticity can hinder the process (Crispo, 2007; Murren et al., 2015). Hypothetically, the cost of plasticity may lead to constitutive divergent expression of genes across diverse habitats and selection can reinforce the variability when it is heritable and gene flow is weak. Our reciprocal transplantation framework allows to identify the constitutive and plastic component of expression divergence and evaluate factors that might constrain and/or facilitate this divergence in the context of ecological variability. For instance, we found *CPK13* (*P.hallii* ortholog of *A. thaliana* AT3G51850.1 *Calcium dependent Protein Kinase 13, PhHAL.2G319400*) and *PHS1* (*P. hallii* ortholog of *A. thaliana* AT5G23720.1-*Propyzamide-Hypersensitive1, PhHAL.5G392700*) were constitutively highly expressed in leaf of both inland genetic clusters compared to the coastal cluster. These genes were previously detected as *cis* eQTL candidates by Lovell et al (2019) (Supplemental Table S7). CPK13 controls stomatal behavior by regulating the expression of guard cell potassium channels (KAT1 and 2), whereas PHS1 controls stomatal behavior via abscisic acid (ABA) signaling. The constitutive expression of CPK13 and PHS1 may provide fitness benefits for *P. hallii* that are native to water-deficit inland habitats, but it may have a trade-off in wet coastal habitats due to the high cost of expression.

### Recombination landscape delineates excess of constitutive expression divergence in low recombining regions

Recombination can generate novel allelic combination by bringing together multiple beneficial alleles or decoupling them from deleterious alleles. Recombination rate can vary between different species, genetic clusters within species, or even along chromosomes. Grass species such as rye (Schreiber et al, 2022) and barley (Dreissig et al, 2019) show striking variation of recombination rate along the length of their chromosomes. In the same line of these reports, *P. hallii* exhibits remarkable variation in the recombination landscape across chromosomes, leaving large parts of the chromosome near pericentromeric regions as mostly non-recombining (Figure 1C; Supplementary Figure S2). The genomic landscape of recombination can play a key role in shaping the pattern of diversity along the genome by altering the efficacy of selection. Indeed, a different pattern of expression divergence across the recombination landscape was observed. We detected that putative divergently expressed genes (both G category genes detected in factorial model and CDE genes detected by outlier test) were enriched in genomic regions with lower recombination rates. However, plastic genes (GxE) were significantly depleted in these regions. Excessive constitutive expression divergence associated with these regions could be an indicator of maladaptive expression divergence; it could arise due to inefficient purifying selection. Expanding our test for positive selection on protein evolution including genes in the low-recombining regions detected no excess of positive selection (p-value< 0.78) further supports the idea.

### Association of expression divergence with sequence evolution

Different trends of association between protein evolutionary rate and expression divergence were observed among different model systems and taxa (Hodgins et al., 2016; Liao and Zhang, 2006; Moyers et al,2013; Nuzhdin et al.,2004; Renaut et al.,2012). We found significant elevated relative (F_ST_) and absolute divergence (D_XY_) among genetic clusters and faster protein evolution (dN/dS) for CDE genes relative to a genomic background of NDE genes (expressed but lacking expression divergence). Increased fixation rates due to positive selection on CDE genes, reduced purifying selection on these genes in at least one genetic cluster, or both might lead to this relationship. However, we did not observe an excess of CDE genes under positive selection. Therefore, we reason that many CDE genes might have evolved under relaxed purifying selection. Alternatively, the relatively deep divergence time between *P. virgatum* and *P. hallii* (∼6.6 mya) may result in lower power for detecting selection events across *P*. *hallii* lineages.

Studies in both animal and plant systems have detected that gene expression level was associated with sequence evolution and genes with higher expression level evolved more slowly (Hodgins et al., 2016, Duret and Mouchiroud, 2000). This could be due to the fact that highly expressed genes tend to interact with a wide range of partners and therefore less likely to tolerate mutation (Duret and Mouchiroud, 2000) or could be under purifying selection due to fitness loss from protein misfolding as a consequence of mutation (Wu et al., 2022). We similarly detected a negative correlation between mean gene expression and the rate of protein evolution (dN/dS). Moreover, we detected that the mean expression level is lower for CDE genes compared to NDE genes. It is possible that moderately expressed genes which were less ubiquitous and hence had fewer pleiotropic effect might have evolved population specific expression patterns that provides fitness benefits in native habitats. Moreover, the cost of plasticity, trade-offs in foreign environments, or lack of gene flow could allow selection to reinforce the difference in expression among different genetic clusters and eventually lead to constitutive divergence. It is likely that the very diverse habitats of inland and coastal *P. hallii* have limited gene flow (even in sympatric habitats), along with extensive inbreeding, might have contributed to the abundance of constitutive divergence.

### Enrichment of cis-eQTLs for putatively divergent gene expression

Cis- and trans-acting regulatory architecture can contribute to expression variation. However, the importance of these molecular mechanisms differ by study system: experiments in yeast, flies and Arabidopsis reported cis-variants accounted for a greater proportion of inter-species gene expression variation (Ronald et al., 2005; Wittkopp et al., 2008; Tirosh et al., 2009) while in maize and *P. hallii cis*-variants accounted for a greater proportion in intra-species expression divergence (Schadt et al., 2003; Guo et al., 2004; Springer and Stupar, 2007, Lovell et al., 2016; Lovell et al., 2018). In this study, we observed an enrichment of both cis- and cis-GXE (*cis*-x drought) eQTLs detected by Lovell et al (2018) in our putatively divergent expression category detected by factorial model (G genes) or CDE genes from Q*_PC_* outlier test. This enrichment of *cis*-eQTL and higher divergence at the promoter region of CDE genes suggest that *cis*-variants might play a greater role in expression divergence of *P. hallii*. *Trans*-variants are expected to be more pleiotropic (Wittkopp, 2007 but also see Lynch and Wagner, 2008) since these interact with a wider range of target genes and hence are likely to have more deleterious effect than *cis*-variants. Hence, it is likely that cis-variants, which exhibit fitness trade-offs between different habitats of *P. hallii*, would be ideal candidates for expression divergence and could be further reinforced by natural selection when gene flow is limited between genetic clusters.

Broadly, reciprocal transplantation experiments integrated with a Q*_PC_* outlier framework allowed us to identify several hundred genes with putatively adaptive constitutive expression divergence but only a weak signal for adaptive plastic divergence. This suggests adaptive plastic responses are rare or were undetected in our experimental context. We observed elevated sequence divergence of promoters and coding sequence of constitutive expression outliers, yet found little evidence for positive selection on these proteins. Furthermore, enrichment of genes with constitutive expression with *cis*-eQTL genes strengthen the hypothesis that *cis*-regulatory divergence as a primary driver of expression variation in *P. hallii.* Together, our study provides a unique look at the evolutionary forces driving patterns of expression polymorphism and divergence in a native grass and identifies outlier candidate genes for future functional genomic studies.

## Methods

### SNP Calling and Filtering

Details for the field collections of *P. hallii* are provided in Palacio-Mejía et al., 2021 and describe collection campaigns from natural field sites distributed across southwest USA (Figure 1A, S1A, and Supplemental Table 1). The geographic range of the *P. hallii* var. *hallii* is larger compared to *P. hallii* var. *filipes* and thus the sample size is greater for the former (75 var. *hallii* and 11 var. *filipes*). *P. hallii* is a strongly selfing species (mean inbreed coefficient 0.89) and field collected material is naturally inbred and highly homozygous (Lowry et al., 2015). Some of these accessions and resequencing data were included in previous studies to investigate genetic variation and population structure (Gould et al. 2018; Lovell 2018). Here, we include 11 newly resequenced accessions (Supplemental Table 1). In order to call Single Nucleotide Polymorphism (SNP) variants we first filtered raw reads for each genotype using BBDuk program from BBTools suite (Bushnell et al. 2017; Bushnell 2020) with a minimum average quality of 20. Next, quality trimmed reads were mapped against the *P.hallii* var. HAL2 reference genome v2 (https://phytozome-next.jgi.doe.gov/info/PhalliiHAL_v2_1) using Burrows-Wheeler Alignment (BWA) mem algorithm (Li and Durbin 2009) with default parameters. We chose the HAL2 reference genome over FIL2 since the HAL2 reference has near complete chromosome level assembly with largest N50 and fewest unassigned scaffolds at chromosome level (Lovell et al, 2019). Moreover, sample sizes for var. *hallii* are much larger compared to var. *filipes*. We further filtered alignments with the samtools software package (Li et al. 2009) with a minimum mapping quality of 20. Filtered alignments were sorted by coordinates, duplicated reads were marked, read groups were assigned for individual genotypes and finally alignments were indexed by Picard tool (v2.20.4) (http://broadinstitute.github.io/picard/) (Toolkit 2018). Alignments around the insertion/deletion regions were masked by the IndelReligner tool and first iteration of SNPs were called by the HaplotypeCaller algorithm from the Genome Analysis Toolkit (GATK) version 3.8.1 (McKenna et al. 2010). Later base recalibration was performed using a set of high-quality variants by the BaseRecalibrator function and the second iteration of variant calling we used the “-- emitRefConfidence GVCF” flag to obtain confidence scores for each site in the genome irrespective of variants or not. Finally, we used GenotypeGVCFs program to collect variant calls and confidences across all individuals and produce genotype calls for each site by setting the “-allSites” flag. We then selected SNP variants and filtered for SNP calling based on the hard filtering recommendation by GATK (https://github.com/tahia/SNP_calling_GATK). SNPs which passed these quality filters were retained as high-quality SNP variants for this diversity panel. Variant sites were further filtered sequentially to keep: i) only biallelic variants, ii) any genotype with DP >=3 was kept else masked as missing, iii) sites kept which have percent of heterozygosity < 3.6 (85^th^ percentile of the distribution), and iv) <20% missing data and finally obtained 23,692,316 high quality biallelic SNPs. Nonvariant sites were separated by VCFtools (Danecek, et al., 2011) and filtered by read depth, quality and percentage of missing individuals (--minDP 20 –minQ 30 –maxmissing 0.8).

### Analysis of population genetic variants and structure

As with previous studies that included the presence of highly divergent groups (e.g. subspecies, ecotypes, etc.), population genetic structure was first assessed hierarchically (Walsh et al. 2017; Napier et al. 2019; Lovell et al. 2021). Specifically, due to the strongly differentiated ecotypes present in our study system, we first identified the broadest genetic population structure using discriminant analysis of principal components (DAPC) in the adegenet (v2.0.1) package (Jombart et al. 2010; Jombart and Ahmed 2011). As this method is not based on common assumptions underlying many population clustering approaches (e.g. Hardy-Weinberg equilibrium and linkage equilibrium), it is a valuable tool for examining broad structural divisions. High quality SNPs were LD pruned (--indep 50 5 2) using plink (Purcell et al. 2007) and annotated by SnpEff tool (Cingolani et al. 2012) with the HAL2 reference genome annotation. From the SNP annotation, synonymous SNPs in coding sequence were selected as putatively neutral variants and were further filtered for minor allele count and missing data (<10% missing data or minor allele count <3). We used a random set of 50,000 these synonymous SNPs for genetic cluster analysis. Prior groups were determined by first transforming the genetic data using principal components analysis (PCA), then the first 10 PCs were used in a k-means algorithm to classify individuals into the broadest two possible groupings aiming to maximize the variation between groups. DAPC was then implemented using 10 retained principal components to provide a description of the genetic clusters (i.e. the two ecotypes).

After classifying individuals into two broad genetic groupings consistent with the ecotypes, we evaluated the genetic structure of and potential admixture between individuals belonging to the more well-represented *hallii* group using the Bayesian clustering algorithm implemented in STRUCTURE (v2.3.4) (Pritchard et al. 2000; Falush et al. 2003). Specifically, the same set of loci were used, but we subset the individuals to those with a DAPC posterior probability of greater than 0.95 for belonging to the *hallii* genetic subpopulation. Within STRUCTURE, we utilized an admixture model with correlated allele frequencies and no prior information. The analysis entailed 20,000 burn-in steps and 100,000 replicates of 1–8 genotypic groups (K), each of which was run 10 times. Using STRUCTURE HARVESTER (Earl and vonHoldt 2012), the resulting output was compiled and evaluated for the optimal K value based on ΔK (Evanno et al. 2005). One accession (COL) which demonstrated small degree of admixture between two inland genetic clusters was removed for further analyses.

### Estimation of population-scaled recombination rate

We used the high-quality imputed SNP calls of three different genetic clusters to estimate recombination rate (ρ/kb, ρ = 4Ne × re, where Ne is the effective population size and re is the effective recombination rate per generation in a population). To account for different sample size of each genetic cluster we randomly subsample individuals from each population (n=11) for this study. First, SNPs were filtered with the following criteria: i) all heterozygous call were masked as missing, ii) SNPs with less than 95% missing data were retained, iii) minor allele frequency > 0.1 were retained and iii) thinned by 50 bp distance. Subsequently, SNPs were imputed by Beagle V5.4 (window=5, overlap=2, iterations=30, err=0.001, burnin=10; Browning et al, 2018). The interval program of LDhat package (Auton and McVean, 2007) was used to estimate the ρ/kb for each non-overlapping 5 Mb intervals (theta 0.001, -its 60,000,000 -samp 5000 -bpen 5). We used the stat program of the same package to summarize ρ/kb values and the first 50,000 iterations were removed as burn in. Later, mean recombination was estimated over 1 Mb interval with a step size of 100 Kb. While comparing recombination landscapes across different genetic clusters, ρ values were scaled within each cluster by dividing by the maximum value. The minimum overlapping contiguous regions with elevated recombination rate across genetic clusters has been defined as highly recombined regions (Supplementary Table S2). We used a Recombinant Inbreed Lines (RIL) mapping population generated from inland and coastal references (Haque et al., 2022) to estimate the correlation between recombination rate and cross over evens. The number of crossover events was estimated in 1Mb intervals with a step size of 100 Kb using the *find.breaks* function from “xio” package (Broman et al., 2002).

### Field sites and reciprocal transplantation

We carried out a reciprocal transplant experiment in 2017 at two different field sites: one representing an inland habitat (The Brackenridge Field Laboratory of the University of Texas at Austin, Austin, TX, USA: 30° 15’ 53.928’’ N, 97° 44’ 47.7492’’ W; hereon referred as BFL) and another representing a coastal habitat (USDA Plant Materials Center, Kingsville, TX, USA: 27° 30’ 57.456’’ N, 97° 56’ 34.134’’ W; hereon referred as KINGS) (Supplementary Figure S4A and B). The distance between these inland and coastal field sites is approximately 232 miles (374 km). The two field sites have varied soil composition in terms of available soil nutrients, and soil conductivity. The soil at the coastal site contained more cations and higher electrical conductivity than the soil at the inland site (Supplementary Figure S5A and B). The mean temperatures and at the coastal and inland sites (March –June) were 25.1 °C and 23.1 °C respectively.

All accessions were grown from seed in a greenhouse (14-h days at 500 μE m −2 s −1, 28°C; 10-h nights at 24°C) located at the Brackenridge Field laboratory in Austin Texas. On 8 March, seeds were scarified with sand paper in order to remove the seed coat. Scarified seeds were placed on wet sand in 25 mm deep Petri dishes which were then sealed with parafilm, moved to a greenhouse bench and randomly cycled to standardize environmental conditions. On 15 March, seedlings were transplanted to 2.5-inch pots (SVD-250-BK, T.O. Plastics, Clearwater, MN, USA) filled with soil (60:40 mixture of Promix BX, Premier Tech Horticulture, Riviere-du-Loup, Quebec, Canada; and Turface, Profile Products, Buffalo Grove, IL, USA) and arranged in 32 cell trays. On 25 March, seedlings were thinned to leave one plant per pot and at least six healthy individuals per accession were selected for field transplantation. Plants were further randomized within trays into the field experiment layout. Trays were then randomly cycled in the greenhouse once every week and irrigated as needed.

Our experimental design consisted of 86 *P. hallii* accessions grown at two field locations with a minimum of 3 biological replicates. Field transplantation was performed on 7 April at the inland site and 12 April at the coastal site. Soil at each planting site was tilled to a depth of 6 inches with a roto-tiller and weed barrier fabric (Sunbelt 3.2 oz, Dewitt Company, Sikeston, MO, USA) was applied to the planting area for weed control. Holes large enough to accommodate the transplants and their expected growth were cut in the weed fabric in a honeycomb fashion. Three individuals from each accession were reciprocally transplanted to the field in a randomized design at each field site. The inland site was ten individuals wide and 84 individuals long, and the coastal site was 21 individual wide and 37 individuals long, with a spacing of 0.75 m between individuals. Plants were carefully pulled out from the pots along with soil and sunk directly into the ground. A border row of *P. hallii* surrounded both field sites and was used to control for edge effects.

### Tissue collection for RNA

Tissue collection for RNA and phenotypic measurements took place on 14 June at the inland site and 20 June at the coastal site between 9:30 am and 12:15 pm. Both collection days had mostly clear skies, with mean local temperatures during collection periods of 29.55 °C at the inland site and 30.28 °C at the coastal site. Tissue collection for RNA and relative water content (RWC) was done simultaneously at the time of collection. The tissue collection for RNA was standardized across the genotypes and sites by collecting fully expanded flag leaves from each plant. All plants were at the reproductive stage during the time of tissue collection. In brief, a leaf was excised from the plant and immediately chopped into pieces (∼1-2 cm) in 2 mL Eppendorf tubes loaded with three stainless beads. Samples were flash-frozen in liquid nitrogen and moved to a -80 freezer.

### RWC measurement and statistical analysis

For RWC measurements, two leaves were cut at the ligule and taken from each plant during each sampling period. For particularly small plants, a third leaf was harvested. Fresh weight of leaves was recorded and the cut petiole end of each leaf was then placed into a 15 mL Falcon tube, with roughly 5 mL of water and both leaves were held in place with a cotton ball. Leaves were then stored in a dark cooler while in the field, and transferred to a refrigerator, for a total of ∼ 8 hours to reach full rehydration. The turgid weight was then recorded, and the leaf tissue was subsequently dried in a 65 C oven for 24 hours to record the dry weight. RWC was calculated as (Fresh Weight – Dry Weight) / (Turgid Weight -Dry Weight) x100. We analyzed RWC using a two-way ANOVA model testing for the effect of Genetic clusters (G), field environment (E), and their interaction (GxE). The effect of genetic clusters was tested separately by site using one-way ANOVA and pairwise comparisons between the means of genetic clusters was computed using Tukey HSD.

### RNA extraction and library preparation for sequencing

Leaf tissue was ground in to a fine powder using Geno/Grinder 2010 (SPEX SamplePrep. Metuchen, NJ). RNA extraction was performed according TRIzolTM Reagene user guide (Cat # 15596018). In brief, the ground leaf tissue powder was homogenized in 1 mL of TRIzol reagent and then incubated with 200 µL of Choloroform:Isoamyl alcohol 24:1 (Sigma, C0549) for 10 min on rotator mixer at room temperature followed by centrifugation. The clear supernatant was then transferred to new 1.5 mL tube and added an equal volume (V/V) of isopropanaol to precipitate nucleic acid. Samples were mixed well and the nucleic acid pellet was obtained by centrifugation. The pellet was washed with 75% alcohol, air dried for 10 min at room temperature and suspended in 30 µL of 10 mM Tris-Cl (pH 8.0) to obtain RNA. The RNA samples were treated with DNaseI (AmbionTM, AM2222) to remove contaminating genomic DNA. RNA concentration was measured using Nano-Drop ND-1000 spectrophotometer.

We constructed RNA-seq libraries using a high-throughput 3′-mRNA Tag-Seq 3’-tagseq approach (Meyer et al., 2011; Lohman et al 2017; Weng et al., 2022). Tagseq is widely used for constructing libraries directed at the 3’end of mRNA fragments enriched in a size range of 300-500 base pairs (bp). Briefly, 1 µg of RNA from each sample was incubated at 70 °C to achieve fragmented RNA molecules of desired range. The entire fragmented RNA was used in first strand cDNA synthesis using RNA oligo primer (S-Ill-swMW) and cDNA amplification. Further, 50 ng of purified cDNA was barcoded using Illumina specific barcodes (ILL-BC) and multiplexed using Illumina TruSeq universal adapters (TruSeq-Un). An equal volume of 30 randomly chosen libraries were pooled together and ran on agarose gel to excise the fragment in the 400-500 bp size range. Resulted library pool was loaded on one lane of an Illumina HiSeq-2500 analyzer at the Hudson Alpha genomic sequencing center, Huntsville, AL, USA. We recovered between 5-7 million 100 bp single end (SE100) reads per sample.

### Tag-seq Sequence data processing

In total, we sequenced 508 Tag-seq libraries from samples collected from replicated accessions grown at our inland and Coastal field sites. A few samples were lost due to poor RNA extraction or library construction failure. On average we sequenced 6 libraries from each accession (inland: 249 and Coastal: 259). We used FastQC to evaluate the sequencing quality (Andrews 2017) for various quality matrices such as: average sequence quality, GC-content, k-mer profile and presence of adapter sequences. These matrices were used for finding optimum parameters for quality filtering of raw sequences. We trimmed the first 9 bases (including degenerate bases) from the sequence adapter and homopolymers (polyA or polyT) for base count of >=15mer using Cutadapt (Martin 2011). We allowed reads with an average sequencing quality of >= 20 and retained minimum sequence length of 70 bases. We generated several k-mer profiles with varied k-mer lengths for the *P. hallii* transcriptome and asked what percentage of the total transcriptome can be detected uniquely with that specific k-mer length across the total length of all transcripts. This analysis showed that ∼82% and ∼90% of unique transcripts can be detected by k-mer length of 50 bp and 100 bp, respectively. Therefore, we chose 70 bases as the minimum sequence length to detect approximately 85% of unique transcripts for this genome. Filtered reads were then mapped to *P. hallii* var. *halli*i HAL2 reference genome (v2.0) using BWA-mem algorithm (Li and Durbin 2009). Mapped reads were further filtered using SAMtools and the reads with mapping quality >= 10 were retained. Duplicated alignments were marked by Picard (http://broadinstitute.github.io/picard/). We used Feature counts (Liao et al. 2014) to generate an expression count matrix from filtered and duplication marked alignments using the existing annotated gene models of *P. hallii*. Only first strand-specific reads were considered as a count that has fragment length>= 70 bp (--readExtension3 0 –readExtention5 0 -d 50 -s 1). Sequencing results and mapping statistic are given in the Supplemental Table S13.

### Differential gene expression analysis

We detected gene expression for 18,773 genes from 33,263 annotated gene models of the HAL2 reference genome. To improve DEG (differential gene expression) detection sensitivity, we first removed genes that had low expression by filtering those with an average count < 1 across all 508 samples. We used DESeq2 package in R for DEG analysis (Love et al. 2014). Libraries were first normalized for their size and gene counts were transformed by variance stabilizing transformation (VST) using the fitted dispersion-mean relationship. We used 1000 genes with the highest variance after normalization and transformation in the principal component analysis (PCA) to visualize the global variation in gene expression between samples.

For DEG analysis, we used DEseq2 to fit a factorial model to test for the effect of genetic cluster/origin (G), field environment/transplant site (E) and their interaction. First, the raw counts were filtered, normalized and transformed as described in the earlier section. Each genotype was assigned to one of three genetic clusters by the maximum posterior probability of STRUCURE and DAPC analysis. In brief, we fit negative binomial models for each gene and run likelihood ratio tests (LRT) comparing the full model with reduced model by dropping the factor of interest.

Subsequently, we conducted post hoc analysis on significantly DEGs of this omnibus test to test for G, E and GXE effect of each of the three pairwise comparisons with appropriate contrast matrix by Wald test. Subsequently, we estimated log2 fold change (LFC) of each pairwise contrast separately for each factor (G or E) using the appropriate contrast for scatter plots. To partition the variance components for G, E and G×E, we used the variance Partition package from Bioconductor (Hoffman and Schadt 2016) on voom-transformed data. Variance components were plotted as ternary plots using a customized R script.

In order to estimate the plasticity of each genetic cluster due to transplantation we implemented DAPC function using the adegenet (v2.0.1) package. Linear discriminant function was built by defining native genetic clusters at native sites as separate groups (inland genotypes at inland site and coastal genotypes as coastal site) on the total transcriptome space. DAPC uses higher order PCA to maximize variation between predefined group while minimizing the variance of within group. We measured the plasticity of a focal genotype as the absolute difference between the DAPC value (z-transformed at each genetic cluster level) at its transplant and native sites. We tested the effect of genetic clusters on gene expression plasticity by fitting a linear mixed model with genetic cluster as fixed effect and genotypes as random effects. The mean plasticity of each genetic cluster was presented by the arrow length in Figure 2D.

### Detection of selection on gene expression

We implemented the Q_PC_ method of Joseph et al (2019) to detect natural selection in structured populations using kin relationships determined from genomic data. This method is an extension of the Q_ST_-F_ST_ method where the additive genetic variance (V_A_) is estimated by the orthogonal Principal Components (PCs) of the population kinship matrix. The kinship matrix was constructed and mean-centered using the same set of 50,000 neutral SNPs that were used to construct genetic clusters. Matrix construction and standardization was performed following the method described in Joseph et al (2019). PCs that cumulatively explained the top 25% of variation in the conditional kinship matrix were chosen for *Q*_PC_ testing. Later PCs which explained cumulatively ∼50% of the total variance of the kinship matrix were used as representative of the neutral evolution. We implemented the QPC test on the mean-centered normalized mean of across sites and the difference between sites of each expressed gene. This strategy relied on reaction norm perspective: mean expression is equivalent to studying the main effect of genetic clusters (hereby identified as “constitutive”) whereas the difference is equivalent to the interaction of genetic clusters with planting location (hereby identified as “plastic”). First, Tag-seq libraries were normalized for their library size, were Variance Stabilizing Transformed which uses the fitted dispersion-mean relationship in order to achieve homoscedasticity and then obtained the mean across and difference between sites for each accession. Confidence interval for a given PC for a given trait was estimated as described by Joseph et al (2019). FDR adjusted q-values were calculated using qvalue function in R with an FDR threshold of 0.1 (Storey and Tibshirani 2003; Storey et al., 2004). As mentioned earlier, we observed very low recombination for the heterochromatin regions of the genome compared to euchromatin regions which could alter the expected neutral evolution between these two regions. Therefore, we carried out *Q_PC_* tests in two different approaches: i) Relaxed Mode: where we applied *Q*_PC_ method on all expressed genes (18,773) and ii) Constrained Mode : only on selected genes resided in highly recombined regions (15,735). For the constrained mode we also purged the SNPs which were located in non-recombining regions (7,562 SNPs) and constructed the kinship matrix as described for relaxed mode.

### Orthologs Identification

Orthologs among *Panicum hallii* HAL2 and FIL2 ecotype genome references (HAL2 v2.1: https://phytozome-next.jgi.doe.gov/info/PhalliiHAL_v2_1; FIL2 v3.1: https://phytozome-next.jgi.doe.gov/info/Phallii_v3_1) were identified by OrthoFinder version 2.5.4 applying default parameters except selecting “blast” as the sequence search algorithm. This program uses a heuristic analysis of pairwise sequence similarity to estimate the phylogenetic relationship between genes and infer putative orthologous clusters. We only considered one-to-one ortholog pairs for downstream analysis of sequence evolution to avoid potential complications from paralogous relationships. We also eliminated ortholog pairs with average percent identity below 50% or with greater than 10% difference of protein length in order to exclude misalignment or erroneous orthologous relationship. In this way we detected 19,965 one-to-one ortholog pairs in *P. hallii*. In order to obtain orthologous relationships among the broader *Panicum* lineage, we also considered the K and N subgenomes of allopolyploid *Panicum virgatum* as two additional genomes along with two genomes *of P. hallii* (HAL2 and FIL2) and implement the same OrthoFinder pipeline described above. In *Panicum*, we detected 8,395 one-to-one ortholog pairs considering all pairwise relationships. In order to infer the ancestry of inland and coastal populations, we further identified 7,419 one-to-one orthologs between inland and coastal references of *P. hallii*, two subgenomes of *P. virgatum,* and *Setaria viridis* by implementing the same orthology strategy described above.

### Phylogenetic analysis and ancestry inference

To infer whether the inland or coastal genetic cluster is ancestral, we first aligned 7,419 one-to-one orthologs of *P. hallii*, *P. virgatum* (both N and K subgenomes) and *S. viridis* gene models for individual orthogroup by Clustal Omega (Sievers et al, 2011). Alignments were concatenated to construct the species tree using maximum Likelihood based algorithm, RAxML (version 8.2.12) (Stamatakis 2014) under PROTGAMMA model using species other than *P. hallii* as outgroups with 1000 bootstrap replicates. In order to polarize the tree, we looked at 4969 orthogroups which have at least 3 variant sites per alignment between inland and coastal genetic clusters and for which ancestry can be inferred from outgroups. Subsequently, we estimated the frequency of coastal allele as ancestral for orthogroups with 10,000 bootstrap replicates.

### Molecular Population Genetic Analyses

Our project generated high quality resequencing data from 86 *P. hallii* genotypes for molecular population genetic analyses. To understand global genetic diversity among and between genetic clusters we estimated genome-wide diversity statistics on 20 Kb window (window length=20 Kb, step size=20 Kb) for each pairwise comparison between genetic clusters (F_ST_ and D_XY_) or within each genetic cluster (π). We used VCFtools to estimate mean Weir and Cockerham’s F_ST_. Nucleotide diversity (π) and the absolute measure of divergence between populations (D_XY_) were estimated for the same intervals by pixy tool (Korunes and Samuk, 2021) using all detected sites (both high quality SNP variants and quality filtered nonvariant sites). One-way ANOVA followed by Tukey’s post-hoc tests were performed to test for differences of these statistics between different pair-wise contrasts (or between different genetic clusters for π). Later we also estimated these statistics split by different gene features (CDS, promoter, intron, 5′-UTR, and 3′-UTR) along with complete gene models of one-to-one ortholog pairs which were detected as expressed in our study using HAL2 v2.1 annotation (https://phytozome-next.jgi.doe.gov/info/PhalliiHAL_v2_1).

We determined the pairwise divergence of nonsynonymous and synonymous sites and their ratio in the coding regions of 13,223 one-to-one ortholog genes in *P. hallii* which were detected as expressed in our study using the CODEML function of PAML tool (version 4.9; runmode -2, F3X4 codon frequency). Alignment sites with gaps were removed and ortholog pairs shorter than 50 amino acids were purged. Low divergence at synonymous site may inflate or overestimate the dN/dS ratio therefore ortholog pairs with dS < 0.01 were also excluded. Mann-Whitney two tests were performed to detect for the difference of mean dN/dS between CDE and NDE group. Partial correlation tests (Spearman’s test) for constitutively divergence expression with substitution rate were performed by ‘pcor’ function in R while controlling for mean expression across site. CDE (=1) and NDE (=0) genes were dummy coded to obtain divergence score.

In order to obtain more power to detect site specific positive selection we included two sub-genomes of *Panicum virgatum* and we used site models using CODEML function of PAML (F3X4 codon frequency). An unrooted species tree was provided and dN and dS at each codon across all branches in the tree were estimated. We tested for sites evolving by positive selection (ω=dN/dS is greater than 1) by comparing model M1a (nearly neutral) and model M2a (positive selection) using a likelihood ratio test with twice the difference of the log-likelihood values of the M1a to M2a model Chi-square distributed with 2 degrees of freedom. We used a significance threshold alpha=0.05 and multiple test correction was implemented by Bonferroni method.

## Acknowledgements and Funding Sources

The work (proposal: 10.46936/10.25585/60001016) conducted by the U.S. Department of Energy Joint Genome Institute (https://ror.org/04xm1d337), a DOE Office of Science User Facility, is supported by the Office of Science of the U.S. Department of Energy operated under Contract No. DE-AC02-05CH11231. This project received funding from the National Science Foundation Plant Genome Research Program (IOS-1444533) and by the US Department of Energy, Office of Science, Office of Biological and Environmental Research, Genomic Science Program Grants DE-SC0014156 and DE-SC0021126.

## Author Contributions

G.B.B., T.H., and T.J. planned and designed research. J.B helped manage field experiments and data collection. G.B.B., T.H., and J.N. analyzed data. J.S. supported resequencing efforts. G.B.B., T.H., and T.J. wrote the manuscript, with review and contributions from all authors.

## References

Auton, A., and McVean, G. (2007). Recombination rate estimation in the presence of hotspots. Genome Research 17(8), 1219–1227. doi: 10.1101/gr.6386707.

Broman, K.W., Rowe, L.B., Churchill, G.A., and Paigen, K. (2002). Crossover Interference in the Mouse. Genetics 160(3), 1123–1131. doi: 10.1093/genetics/160.3.1123.

Brooker, R., Brown, L.K., George, T.S., Pakeman, R.J., Palmer, S., Ramsay, L., et al. (2022). Active and adaptive plasticity in a changing climate. Trends in Plant Science 27(7), 717–728. doi: 10.1016/j.tplants.2022.02.004.

Browning, B.L., Zhou, Y., and Browning, S.R. (2018). A One-Penny Imputed Genome from Next-Generation Reference Panels. The American Journal of Human Genetics 103(3), 338–348. doi: https://doi.org/10.1016/j.ajhg.2018.07.015.

Bushnell B. 2020. BBMap. sourceforge. net/projects/bbmap.

Bushnell B, Rood J, Singer E. 2017. BBMerge – Accurate paired shotgun read merging via overlap. PLoS One 12:1–15.

Campbell-Staton, S.C., Velotta, J.P., and Winchell, K.M. (2021). Selection on adaptive and maladaptive gene expression plasticity during thermal adaptation to urban heat islands. Nature Communications 12(1), 6195. doi: 10.1038/s41467-021-26334-4.

Cingolani P, Platts A, Wang LL, Coon M, Nguyen T, Wang L, Land SJ, Lu X, Ruden DM. 2012. A program for annotating and predicting the effects of single nucleotide polymorphisms, SnpEff: SNPs in the genome of Drosophila melanogaster strain w1118; iso-2; iso-3 Fly (Austin). 6:80–92.

Crispo E. 2007. The Baldwin effect and genetic assimilation: revisiting two mechanism of evolutionary change mediated by phenotypic plasticity. Evolution 61:2469–2479.

Dapper, A.L., and Payseur, B.A. (2017). Connecting theory and data to understand recombination rate evolution. Philosophical Transactions of the Royal Society B: Biological Sciences 372(1736), 20160469. doi: doi:10.1098/rstb.2016.0469.

Des Marais DL, Hernandez KM, Juenger TE. 2013. Genotype-by-environment interaction and plasticity: Exploring genomic responses of plants to the abiotic environment. Annu. Rev. Ecol. Evol. Syst. 44:5–29.

Duret, L., and Mouchiroud, D. (2000). Determinants of Substitution Rates in Mammalian Genes: Expression Pattern Affects Selection Intensity but Not Mutation Rate. Molecular Biology and Evolution 17(1), 68–070. doi: 10.1093/oxfordjournals.molbev.a026239.

Dreissig, S., Mascher, M., and Heckmann, S. (2019). Variation in Recombination Rate Is Shaped by Domestication and Environmental Conditions in Barley. Molecular Biology and Evolution 36(9), 2029–2039. doi: 10.1093/molbev/msz141.

Earl DA, von Holdt BM. 2012. STRUCTURE HARVESTER: a website and program for visualizing STRUCTURE output and implementing the Evanno method. Conserv. Genet. Resour. 4:359–361.

Evanno G, Regnaut S, Goudet J. 2005. Detecting the number of clusters of individuals using the software STRUCTURE: A simulation study. Mol. Ecol. 14:2611–2620.

Falush D, Stephens M, Pritchard JK. 2003. Inference of population structure using multilocus genotype data: linked loci and correlated allele frequencies. Genetics 164:1567–1587.

Fay, J.C., and Wittkopp, P.J. (2008). Evaluating the role of natural selection in the evolution of gene regulation. Heredity 100(2), 191–199. doi: 10.1038/sj.hdy.6801000.

Fuess L.E., Weber J.N., den Haan S, Steinel N.C., Shim K.C., Bolnick D.I. 2021. Between-population differences in constitutive and infection-induced gene expression in threespine stickleback. Molecular Ecology 30:6791–6805.

Gilad, Y., Oshlack, A., and Rifkin, S.A. (2006). Natural selection on gene expression. Trends in Genetics 22(8), 456–461. doi: https://doi.org/10.1016/j.tig.2006.06.002.

Gould BA, Chen Y, Lowry DB. 2018. Gene regulatory divergence between locally adapted ecotypes in their native habitats. Mol. Ecol. 27:4174–4188.

Groen, S.C., Ćalić, I., Joly-Lopez, Z., Platts, A.E., Choi, J.Y., Natividad, M., et al. (2020). The strength and pattern of natural selection on gene expression in rice. Nature 578(7796), 572–576. doi: 10.1038/s41586-020-1997-2.

Guo, M., Rupe, M.A., Zinselmeier, C., Habben, J., Bowen, B.A., and Smith, O.S. (2004). Allelic Variation of Gene Expression in Maize Hybrids[W]. The Plant Cell 16(7), 1707–1716. doi: 10.1105/tpc.022087.

Haque, T., Bhaskara, G.B., Yin, J., Bonnette, J., and Juenger, T.E. (2022). Natural variation in growth and leaf ion homeostasis in response to salinity stress in Panicum hallii. Frontiers in Plant Science 13. doi: 10.3389/fpls.2022.1019169.

Hendry, A.P. (2015). Key Questions on the Role of Phenotypic Plasticity in Eco-Evolutionary Dynamics. Journal of Heredity 107(1), 25–41. doi: 10.1093/jhered/esv060.

Hodgins, K.A., Yeaman, S., Nurkowski, K.A., Rieseberg, L.H., and Aitken, S.N. (2016). Expression Divergence Is Correlated with Sequence Evolution but Not Positive Selection in Conifers. Molecular Biology and Evolution 33(6), 1502–1516. doi: 10.1093/molbev/msw032.

Hoffman GE, Schadt EE. 2016. variancePartition: Interpreting drivers of variation in complex gene expression studies. BMC Bioinformatics [Internet] 17:17–22. Available from: http://dx.doi.org/10.1186/s12859-016-1323-z

Ingvarsson, P.K. (2007). Gene Expression and Protein Length Influence Codon Usage and Rates of Sequence Evolution in Populus tremula. Molecular Biology and Evolution 24(3), 836–844. doi: 10.1093/molbev/msl212.

Jombart T, Ahmed I. 2011. adegenet 1.3-1: New tools for the analysis of genome-wide SNP data. Bioinformatics 27:3070–3071.

Jombart T, Devillard S, Balloux F. 2010. Discriminant analysis of principal components: a new method for the analysis of genetically structured populations. BMC Genet. 11:94.

Jones H.G. (2007. Monitoring plant and soil water status: established and novel methods revisited and their relevance to studies of drought tolerance, Journal of Experimental Botany, 58:119–130, https://doi.org/10.1093/jxb/erl118

Josephs EB, Berg JJ, Ross-Ibarra J, Coop G. 2019. Detecting Adaptive Differentiation in Structured Populations with Genomic Data and Common Gardens. Genetics [Internet] 211:989–1004. Available from: https://doi.org/10.1534/genetics.118.301786

Kenkel CD, Matz M V. 2016. Gene expression plasticity as a mechanism of coral adaptation to a variable environment. Nat. Ecol. Evol. [Internet] 1:14. Available from: www.nature.com/natecolevol

Khasanova, A., Lovell, J. T., Bonnette, J., Weng, X., Jenkins, J., Yoshinaga, Y., et al. (2019). The genetic architecture of shoot and root trait divergence between mesic and xeric ecotypes of a perennial grass. Front. Plant Sci. 10. doi: 10.3389/fpls.2019.00366

Korunes, K.L. and Samuk, K. (2021), pixy: Unbiased estimation of nucleotide diversity and divergence in the presence of missing data. Mol Ecol Resour, 21: 1359–1368. https://doi.org/10.1111/1755-0998.13326

Li, A., Li, L., Wang, W., and Zhang, G. (2019). Evolutionary trade-offs between baseline and plastic gene expression in two congeneric oyster species. Biology Letters 15(6), 20190202. doi: doi:10.1098/rsbl.2019.0202.

Li H, Durbin R. 2009. Fast and accurate short read alignment with Burrows-Wheeler transform. Bioinformatics 25:1754–1760.

Li H, Handsaker B, Wysoker A, Fennell T, Ruan J, Homer N, Marth G, Abecasis G, Durbin R. 2009. The Sequence Alignment/Map format and SAMtools. Bioinformatics 25:2078–2079.

Li, J., and Burmeister, M. (2005). Genetical genomics: combining genetics with gene expression analysis. Human Molecular Genetics 14(suppl_2), R163–R169. doi: 10.1093/hmg/ddi267.

Liao Y, Smyth GK, Shi W. 2014. FeatureCounts: An efficient general purpose program for assigning sequence reads to genomic features. Bioinformatics 30:923–930.

Liao, B.-Y., and Zhang, J. (2006). Low Rates of Expression Profile Divergence in Highly Expressed Genes and Tissue-Specific Genes During Mammalian Evolution. Molecular Biology and Evolution 23(6), 1119–1128. doi: 10.1093/molbev/msj119.

Lohman, B.K., Stutz, W.E., and Bolnick, D.I. (2017). Gene expression stasis and plasticity following migration into a foreign environment. Molecular Ecology 26(18), 4657–4670. doi: https://doi.org/10.1111/mec.14234.

Love MI, Huber W, Anders S. 2014. Moderated estimation of fold change and dispersion for RNA-seq data with DESeq2. Genome Biol. 15:1–21.

Lovell, J.T., Jenkins, J., Lowry, D.B., Mamidi, S., Sreedasyam, A., Weng, X., et al. (2018). The genomic landscape of molecular responses to natural drought stress in Panicum hallii. Nature Communications 9 (1), 5213. doi: 10.1038/s41467-018-07669-x.

Lovell, J.T., Schwartz, S., Lowry, D.B., Shakirov, E.V., Bonnette, J.E., Weng, X., et al. (2016). Drought responsive gene expression regulatory divergence between upland and lowland ecotypes of a perennial C4 grass. Genome Research. doi: 10.1101/gr.198135.115.

Lovell J.T., MacQueen A.H., Mamidi S, Bonnette J, Jenkins J, Napier J.D., Sreedasyam A, Healey A, Session A, Shu S, et al. 2021. Genomic mechanisms of climate adaptation in polyploid bioenergy switchgrass. Nature 590:438–444.

Lowry D.B., Purmal CT, Juenger TE. 2013. A population genetic transect of Panicum hallii (Poaceae). Am. J. Bot. 100:592–601.

Lowry D.B., Hernandez K, Taylor S.H., Meyer E, Logan T.L., Barry K.W., Chapman J.A., Rokhsar D.S., Schmutz J, Juenger T.E. 2015. The genetics of divergence and reproductive isolation between ecotypes of Panicum hallii. New Phytologist 205:402–414.

Lynch, V.J., and Wagner, G.P. (2008). Resurrecting the role of transcription factor change in developmental evolution. Evolution 62 (9), 2131–2154. doi: https://doi.org/10.1111/j.1558-5646.2008.00440.x.

Martin M. 2011. &Xwdgdsw Uhpryhv Dgdswhu Vhtxhqfhv Iurp Kljk Wkurxjksxw Vhtxhqflqj Uhdgv. EMBnet.journal 17:10–12.

Mäkinen, H., Sävilammi, T., Papakostas, S., Leder, E., Vøllestad, L.A., and Primmer, C.R. (2017). Modularity Facilitates Flexible Tuning of Plastic and Evolutionary Gene Expression Responses during Early Divergence. Genome Biology and Evolution 10(1), 77–93. doi: 10.1093/gbe/evx278.

McKenna A, Hanna M, Banks E, Sivachenko A, Cibulskis K, Kernytsky A, Garimella K, Altshuler D, Gabriel S, Daly M. 2010. The Genome Analysis Toolkit: a MapReduce framework for analyzing next-generation DNA sequencing data. Genome Res. 20:1297–1303.

Meiklejohn, C.D., Coolon, J.D., Hartl, D.L., and Wittkopp, P.J. (2014). The roles of cis- and trans-regulation in the evolution of regulatory incompatibilities and sexually dimorphic gene expression. Genome Research 24(1), 84–95. doi: 10.1101/gr.156414.113.

Metzger, B.P.H., Wittkopp, P.J., and Coolon, J.D. (2017). Evolutionary Dynamics of Regulatory Changes Underlying Gene Expression Divergence among Saccharomyces Species. Genome Biology and Evolution 9(4), 843–854. doi: 10.1093/gbe/evx035.

Meyer E, Aglyamova G.V., Matz M.V., Profiling gene expression responses of coral larvae (Acropora millepora) to elevated temperature and settlement inducers using a novel RNA-Seq procedure. Mol Ecol. 2011 Sep;20(17):3599–616. doi: 10.1111/j.1365-294X.2011.05205.x. Epub 2011 Jul 29. PMID: 21801258.

Moyers, B.T., and Rieseberg, L.H. (2013). Divergence in Gene Expression Is Uncoupled from Divergence in Coding Sequence in a Secondarily Woody Sunflower. International Journal of Plant Sciences 174(7), 1079–1089. doi: 10.1086/671197.

Murren, C.J., Auld, J.R., Callahan, H., Ghalambor, C.K., Handelsman, C.A., Heskel, M.A., et al. (2015). Constraints on the evolution of phenotypic plasticity: limits and costs of phenotype and plasticity. Heredity 115(4), 293–301. doi: 10.1038/hdy.2015.8.

Napier J.D., de Lafontaine G, Heath K.D., Hu F.S. 2019. Rethinking long-term vegetation dynamics: multiple glacial refugia and local expansion of a species complex. Ecography 42:1056–1067.

Necsulea, A., and Kaessmann, H. (2014). Evolutionary dynamics of coding and non-coding transcriptomes. Nature Reviews Genetics 15(11), 734–748. doi: 10.1038/nrg3802.

Nicotra, A.B., Atkin, O.K., Bonser, S.P., Davidson, A.M., Finnegan, E.J., Mathesius, U., et al. (2010). Plant phenotypic plasticity in a changing climate. Trends in Plant Science 15(12), 684–692. doi: 10.1016/j.tplants.2010.09.008.

Nourmohammad, A., Rambeau, J., Held, T., Kovacova, V., Berg, J., and Lässig, M. (2017). Adaptive Evolution of Gene Expression in Drosophila. Cell Reports 20(6), 1385–1395. doi: https://doi.org/10.1016/j.celrep.2017.07.033.

Nuzhdin, S.V., Wayne, M.L., Harmon, K.L., and McIntyre, L.M. (2004). Common Pattern of Evolution of Gene Expression Level and Protein Sequence in Drosophila. Molecular Biology and Evolution 21(7), 1308–1317. doi: 10.1093/molbev/msh128.

Palacio-Mejía JD, Grabowski PP, Ortiz EM, Silva-Arias GA, Haque T, Des Marais DL, Bonnette J, Lowry DB, Juenger TE. 2021. Geographic patterns of genomic diversity and structure in the C4 grass Panicum hallii across its natural distribution. AoB Plants 13.

Pritchard JK, Stephens M, Donnelly P. 2000. Inference of population structure using multilocus genotype data. Genetics 155:945–959.

Price, P.D., Palmer Droguett, D.H., Taylor, J.A., Kim, D.W., Place, E.S., Rogers, T.F., et al. (2022). Detecting signatures of selection on gene expression. Nature Ecology & Evolution 6(7), 1035–1045. doi: 10.1038/s41559-022-01761-8.

Prud’homme, B., Gompel, N., and Carroll, S.B. (2007). Emerging principles of regulatory evolution. Proceedings of the National Academy of Sciences 104(suppl_1), 8605–8612. doi: doi:10.1073/pnas.0700488104

Purcell S, Neale B, Todd-Brown K, Thomas L, Ferreira MAR, Bender D, Maller J, Sklar P, De Bakker PIW, Daly MJ, et al. 2007. PLINK: A tool set for whole-genome association and population-based linkage analyses. Am. J. Hum. Genet. 81:559–575.

Renaut, S., Grassa, C.J., Moyers, B.T., Kane, N.C., and Rieseberg, L.H. (2012). The Population Genomics of Sunflowers and Genomic Determinants of Protein Evolution Revealed by RNAseq. Biology 1(3), 575–596.

Romero, I.G., Ruvinsky, I., and Gilad, Y. (2012). Comparative studies of gene expression and the evolution of gene regulation. Nature Reviews Genetics 13(7), 505–516. doi: 10.1038/nrg3229.

Ronald, J., Brem, R.B., Whittle, J., and Kruglyak, L. (2005). Local Regulatory Variation in Saccharomyces cerevisiae. PLOS Genetics 1(2), e25. doi: 10.1371/journal.pgen.0010025.

Schadt, E.E., Monks, S.A., Drake, T.A., Lusis, A.J., Che, N., Colinayo, V., et al. (2003). Genetics of gene expression surveyed in maize, mouse and man. Nature 422(6929), 297–302. doi: 10.1038/nature01434.

Schaefke, B., Emerson, J.J., Wang, T.-Y., Lu, M.-Y.J., Hsieh, L.-C., and Li, W.-H. (2013). Inheritance of Gene Expression Level and Selective Constraints on Trans-and Cis-Regulatory Changes in Yeast. Molecular Biology and Evolution 30(9), 2121–2133. doi: 10.1093/molbev/mst114.

Schreiber, M., Gao, Y., Koch, N., Fuchs, J., Heckmann, S., Himmelbach, A., et al. (2022). Recombination Landscape Divergence Between Populations is Marked by Larger Low-Recombining Regions in Domesticated Rye. Molecular Biology and Evolution 39(6). doi: 10.1093/molbev/msac131.

Shi, X., Ng, D.W.K., Zhang, C., Comai, L., Ye, W., and Jeffrey Chen, Z. (2012). Cis- and trans-regulatory divergence between progenitor species determines gene-expression novelty in Arabidopsis allopolyploids. Nature Communications 3(1), 950. doi: 10.1038/ncomms1954.

Sievers, F., Wilm, A., Dineen, D., Gibson, T.J., Karplus, K., Li, W., et al. (2011). Fast, scalable generation of high-quality protein multiple sequence alignments using Clustal Omega. Molecular Systems Biology 7(1), 539.

Slotte, T., Bataillon, T., Hansen, T.T., St. Onge, K., Wright, S.I., and Schierup, M.H. (2011). Genomic Determinants of Protein Evolution and Polymorphism in Arabidopsis. Genome Biology and Evolution 3, 1210–1219. doi: 10.1093/gbe/evr094.

Stamatakis A. 2014. RAxML version 8: A tool for phylogenetic analysis and post-analysis of large phylogenies. Bioinformatics 30:1312–1313.

Springer, N.M., and Stupar, R.M. (2007). Allelic variation and heterosis in maize: How do two halves make more than a whole? Genome Research 17(3), 264–275. doi: 10.1101/gr.5347007.

Staubach, F., Teschke, M., Voolstra, C.R., Wolf, J.B.W., and Tautz, D. (2010). A test of the neutral model of expression changes in natural populations of house mouse subspecies. Evolution 64(2), 549–560. doi: https://doi.org/10.1111/j.1558-5646.2009.00818.x.

Stern, D.L., and Orgogozo, V. (2008). The loci of evolution: How predictable is genetic evolution? Evolution 62(9), 2155–2177. doi: 10.1111/j.1558-5646.2008.00450.x.

Storey J.D., Taylor J.E., Siegmund D. 2004. Strong control, conservative point estimation and simultaneous conservative consistency of false discovery rates: A unified approach. J. R. Stat. Soc. Ser. B Stat. Methodol. 66:187–205.

Storey J.D., Tibshirani R. 2003. Statistical significance for genomewide studies. Proc. Natl. Acad. Sci. U. S. A. 100:9440–9445.

Tirosh, I., Reikhav, S., Levy, A.A., and Barkai, N. (2009). A Yeast Hybrid Provides Insight into the Evolution of Gene Expression Regulation. Science 324(5927), 659–662. doi: doi:10.1126/science.1169766.

Vande Zande, P., Hill, M.S., and Wittkopp, P.J. (2022). Pleiotropic effects of trans-regulatory mutations on fitness and gene expression. Science 377(6601), 105–109. doi: doi:10.1126/science.abj7185.

Walsh R, Thomson K.L., Ware J.S., Funke B.H., Woodley J, McGuire K.J., Mazzarotto F, Blair E, Seller A, Taylor J.C., et al. 2017. Reassessment of Mendelian gene pathogenicity using 7,855 cardiomyopathy cases and 60,706 reference samples. Genetics in Medicine 19:192–203.

Weng X, Juenger T.E. 2022. A High-Throughput 3’-Tag RNA Sequencing for Large-Scale Time-Series Transcriptome Studies. Methods Mol Biol. 2398:151–172. doi: 10.1007/978-1-0716-1912-4_13. PMID: 34674175.

Whitehead A, Crawford D.L. 2006a. Variation within and among species in gene expression: Raw material for evolution. Mol. Ecol. 15:1197–1211.

Whitehead A, Crawford D.L. 2006b. Neutral and adaptive variation in gene expression. Proc. Natl. Acad. Sci. U. S. A. 103:5425–5430.

Wittkopp, P.J. (2007). Variable gene expression in eukaryotes: a network perspective. Journal of Experimental Biology 210(9), 1567–1575. doi: 10.1242/jeb.002592.

Wittkopp, P.J., Haerum, B.K., and Clark, A.G. (2008). Regulatory changes underlying expression differences within and between Drosophila species. Nature Genetics 40(3), 346–350. doi: 10.1038/ng.77.

Wu, Z., Cai, X., Zhang, X., Liu, Y., Tian, G.-b., Yang, J.-R., et al. (2022). Expression level is a major modifier of the fitness landscape of a protein coding gene. Nature Ecology & Evolution 6(1), 103–115. doi: 10.1038/s41559-021-01578-x

